# CNCC: An analysis tool to determine genome-wide DNA break end structure at single-nucleotide resolution

**DOI:** 10.1101/611756

**Authors:** Karol Szlachta, Heather M Raimer, Laurey D. Comeau, Yuh-Hwa Wang

**Author notes:** These authors contributed equally to this work (Karol Szlachta, Heather M Raimer). Correspondence to Yuh-Hwa Wang: Department of Biochemistry and Molecular Genetics, University of Virginia, Charlottesville, Virginia, 22903-0733, USA.

## Abstract

DNA double-stranded breaks (DSBs) are potentially deleterious events in a cell. The end structures (blunt, 3’- and 5’-overhangs) at sites of double-stranded breaks contribute to the fate of their repair and provide critical information for consequences of the damage. Here, we describe the use of a coverage-normalized cross correlation analysis (CNCC) to process high-precision genome-wide break mapping data, and determine genome-wide break end structure distributions at single-nucleotide resolution. For the first time, on a genome-wide scale, our analysis revealed the increase in the 5’ to 3’ end resection following etoposide treatment, and the global progression of the resection due to the removal of DNA topoisomerase II cleavage complexes. Further, our method distinguished the change in the pattern of DSB end structure with increasing doses of the drug. The ability of this method to determine DNA break end structures without *a priori* knowledge of break sequences or genomic position should have broad applications in understanding genome instability.

## Introduction

DNA double-stranded breaks (DSBs) are one of the most dangerous types of damage that occur in cells, and when unrepaired or illegitimately repaired, DSBs can be cytotoxic or cause genome instability. Multiple pathways to repair these lesions exist, and DNA end structure at the site of the break is one factor determining which of these pathways is used for repair [1-3]. Homologous recombination (HR) is dependent on homologous sequences to act as a template for repairing the DSB, and requires extensive resection of the ends on both sides of the DSB. Meanwhile, non-homologous end joining (NHEJ) canonically ligates the two DNA strands with limited end-processing, resulting in potentially error-prone ligation products. When a wide range of endogenous and exogenous conditions generate DSBs, different end structures can be produced and then affect the repair efficiency, timing, kinetics, and accuracy of the break sites [4, 5].

The importance of understanding DSBs and their repair prompted a recent eruption of experimental techniques to precisely map/sequence DSBs [6, 7]. Each DSB generates two distinct DNA ends and these methods captured these ends which encode the genomic location of the break. Existing studies have primarily used this DSB location data to evaluate breaks occurring at specific loci or subsets of the genome, as opposed to identifying the global patterns of break end structures in the highly detailed data that their techniques have generated. We noticed that the resulting break data also provides coverage information on both positive and negative strands for each individual break in the entire genome. Therefore, using a coverage-normalized cross correlation (CNCC) between the coverage on the positive and negative strands, the type of end structures at the breaks can be revealed at a single-nucleotide resolution, and the genome-wide composition of DSB end structures can be retrieved from data generated by any given technique. Previous methods that have studied the impact of DSB end structures have primarily relied upon either a few specific loci or DNA fragments in an *in vitro* extract experiment. Therefore, a gap is present in understanding the global impacts of end structures and the repair that follows.

Here, we present the utility of our CNCC method, on a genome-wide level, to distinguish the three major DNA end structures: blunt-ended, 3’-overhangs, and 5’-overhangs, using multiple restriction enzymes. Additionally, we tested the sensitivity of our method to detect an end structure that only represents a small fraction of the acquired break data. Finally, we demonstrate the global endogenous DSB end structure landscape of a non-malignant cell line and the subsequent change upon treatment with etoposide, a chemotherapeutic drug. This change lead to an increase in 5’ to 3’ end resection products following treatment and was dose dependent. Overall, our analysis can both provide a global view of what type of end structures are occurring in cells upon DNA damage, and can be used to assess trends of damage types resulting from different drug treatments and between different regions of the genome.

## Results

### Overview of End-Structure Determination from DSB Mapping Data

Each DSB generates two distinct DNA ends. Those ends can be captured by DNA break mapping/sequencing methods as two distinct DNA fragments, and provide coverage information on the positive and negative strands for individual breaks. This information can then be used to reveal the end structure of broken ends. In the genome-wide break mapping protocol that we employed (see “Materials and Methods”), the ligation of the P5 adaptor captures each of the DSB ends; this allows for the identification of the DSB-proximal nucleotide as the most 5’ nt of read 1 of the sequenced pair for each side of the break (**Fig. S1**). To analyze the types of break end structures at a genome-wide scale, we developed an unbiased analysis approach that uses the profiles of single-nucleotide resolution breaks on the positive and negative strands to calculate the most abundant relative shifts between these two groups of signals (we named “Coverage Normalized Cross Correlation, CNCC”). These relative shifts reveal the genome-wide composition of DSB end types at single-nucleotide resolution, without the explicit knowledge of which reads come from a single DSB site or of sequence composition at DSB sites. CNCC is a correlation between two different signals calculated for a range of relative shifts between them and normalized by the total break coverage of a given sample. Therefore, in our analysis, the relative distance between a pair of genomic positions on positive and negative strand codes what type of DSB end-structure was generated (**Fig. 1a**).

**Fig. 1.**
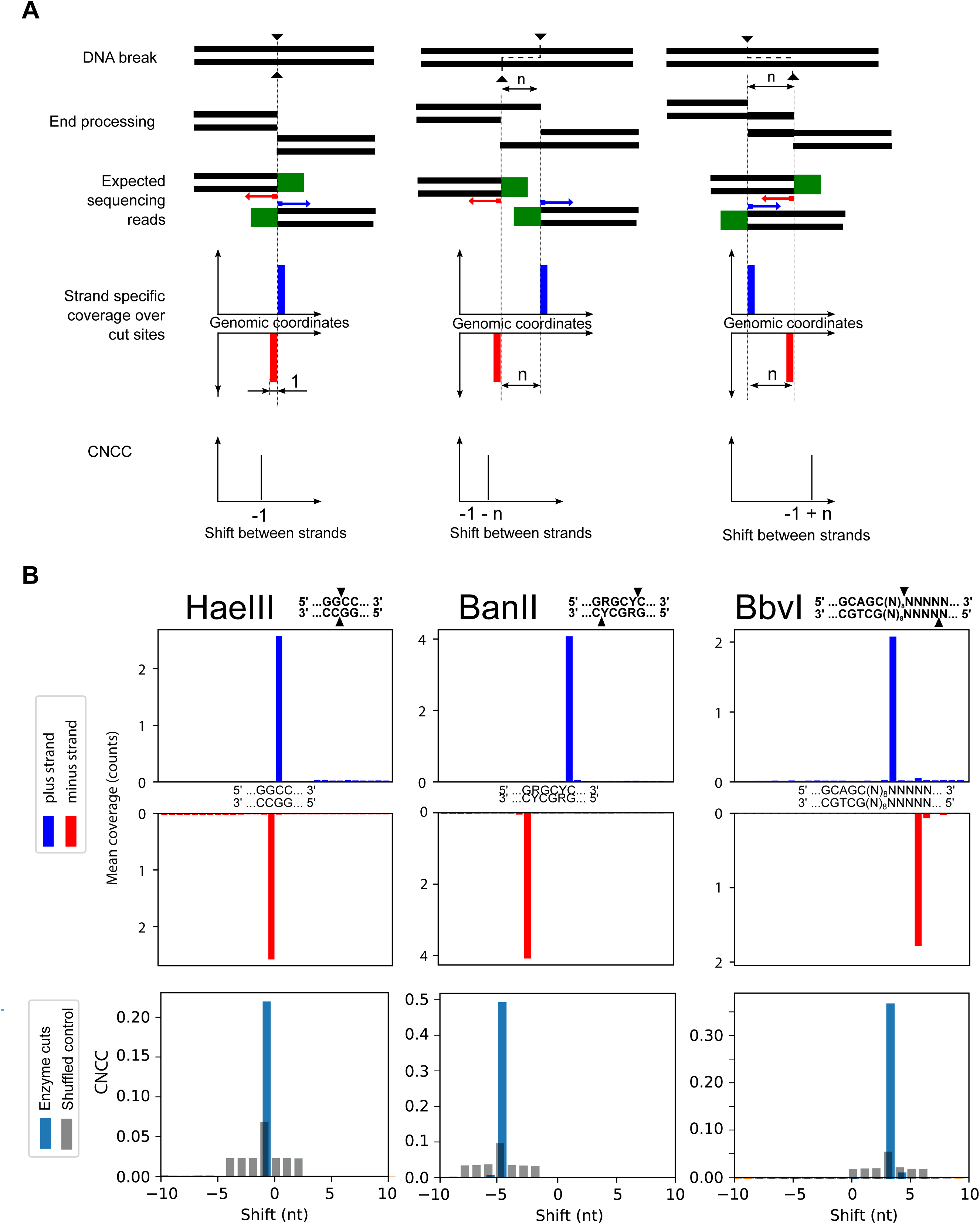
Determination of DSB end structures by CNCC. **(a)** Outline of the CNCC analysis for three types of breaks: blunt end (left), 3’ end overhang (middle) or 5’ end overhang (right). Each break produces two DNA ends and each of those are processed (Materials and Methods), ligated to a sequencing adaptor (green rectangles), and sequenced. These reads result in distinctive patterns of coverage (fourth row). Genome-wide CNCC between coverage on positive and negative strands reveal a shift that is characteristic for the type of DSB end structure (fifth row). (**b)** CNCC analysis of DNA breaks caused by HaeIII, BanII and BbvI restriction enzyme cleavage (left, middle, and right column, respectively). The mean read coverage over enzyme motifs presents precise location of reads (top two rows), and genome-wide CNCC spikes (bottom row, blue) at −1 for HaeIII, −5 for BanII, and +3 for BbvI exactly reflect the expected end structures, with shuffled controls in gray.

### Validation of CNCC End-Structure Determination

To determine the ability of our approach to distinguish different end structures, we evaluated the DNA breaks generated by restriction enzyme cleavage. Digestion of isolated genomic DNA from non-malignant GM13069 lymphoblasts by BanII, BbvI, and HaeIII enzymes respectively produces 3’ overhangs, 5’ overhangs and blunt-end breaks. Break mapping and sequencing of the enzyme-digested samples resulted in 2.0 M, 2.3 M, and 3.9 M enzyme cut sites with reads, and with reads at cut sites accounting for 90%, 61%, and 97% of total reads, respectively (**Table S1**). Each library was sequenced to at least 15 million reads (**Table S2**), and the mean coverage over cut-site regions showed a clear enrichment at cut sites with barely visible background (**Fig. 1b**, top two rows). The genome-wide CNCC analysis, using all mapped DNA ends for each enzyme-digested sample (**Table S2**), demonstrated that restriction enzyme shifts of CNCC (**Fig. 1b**, bottom) spike at −1 for HaeIII, −5 for BanII, and +3 for BbvI precisely corresponded to the expected DNA end types: blunt end, 4-nt 3’ overhang, and 4-nt 5’ overhang, accordingly. Additionally, comparing technical replicates of the BbvI digestion experiment, showed near perfect reproducibility of both genomic coverage and restriction enzyme shift spikes of CNCC between experiments (**Fig. S2**). To further validate our method, we analyzed published data from an EcoRV digestion in HeLa cells [8], and detected an enzyme spike of CNCC at −1 shift which corresponded exactly with the expected blunt end structure generated by EcoRV (**Fig. S3**). Altogether, these results suggest that our CNCC analysis is both robust in its ability to distinguish different end-structures and is highly reproducible.

While validating end structure detection by CNCC, determining the appropriate control for this analysis was also investigated. To establish a meaningful control, first a simple random shuffle was implemented; however, considering that the human genome is on the order of 3×10^9^ nt, performing a simple random shuffle over this large region produced nearly nonexistent background levels. This comes both from the size of the shuffling region being so large and the disruption of break cluster sites by the unrestrained random shuffle. Therefore, to create a more stringent control, we introduced much smaller perturbations to the signal to conserve the overall composition of the break intensities and clustering. First, we shuffled single nt coverage intensities only between regions that originally had non-zero coverage, therefore maintaining break clustering. Second, to each genomic position with non-zero break coverage, we added a random value to wiggle the position in order to control the size of the region that the signal is shuffled in. A series of different wiggle magnitudes were tested on the HaeIII data to establish which would give the most meaningful control (**Fig. S4a**). We determined that the magnitude of the wiggle directly corresponds to the maximum spike range the control is needed for. This allows a meaningful shuffle and also minimizes the size of the perturbation to obtain an appropriately stringent control. For the restriction enzyme digestion experiments, because the maximum spike range to be controlled for is 1, we then used a wiggle ranging from +2 to −2 (0, no shift, excluded) to generate the shuffled control **(Fig. 1b**, bottom**, Fig. S2 and S3)**.

### Testing CNCC Sensitivity

To assess the sensitivity of CNCC analysis to detect the end structure of DNA breaks present at a low fraction of the total breaks, we processed published break mapping data from a limited AsiSI digestion in U2OS cells [8]. In this system, only a fraction of the total reads (0.03%, ∼30,000 reads) were generated from the AsiSI digestion, compared to nearly 61-97% of reads attributed to enzyme cleavage in our previous experiments (**Fig. 1b**). AsiSI digestion produces a 2-nt 3’ overhang DNA end structure, which corresponds to a shift spike for NCC at −3, and our analysis indeed detected this spike (**Fig. 2a**, top). When all reads located at AsiSI cut sites were found and removed, we saw a concordant drop in the spike of CNCC at −3, without altering any other spikes (**Fig. 2a**, bottom). Next, we calculated CNCC while randomly incrementally masking 10% of cut sites, and performing 100 iterations for each 10% increment of random masking. Results showed that while the shuffled control level remained unchanged (grey), a gradual, linear decrease of CNCC for the −3 spike was observed (blue) (**Fig. 2b**). This exercise clearly demonstrates the high sensitivity of CNCC to detect a break species that exist only as a minimal contributor to total break signal and further detect even small changes in the composition of the DNA break end structure within a larger pool of break signals.

**Fig. 2.**
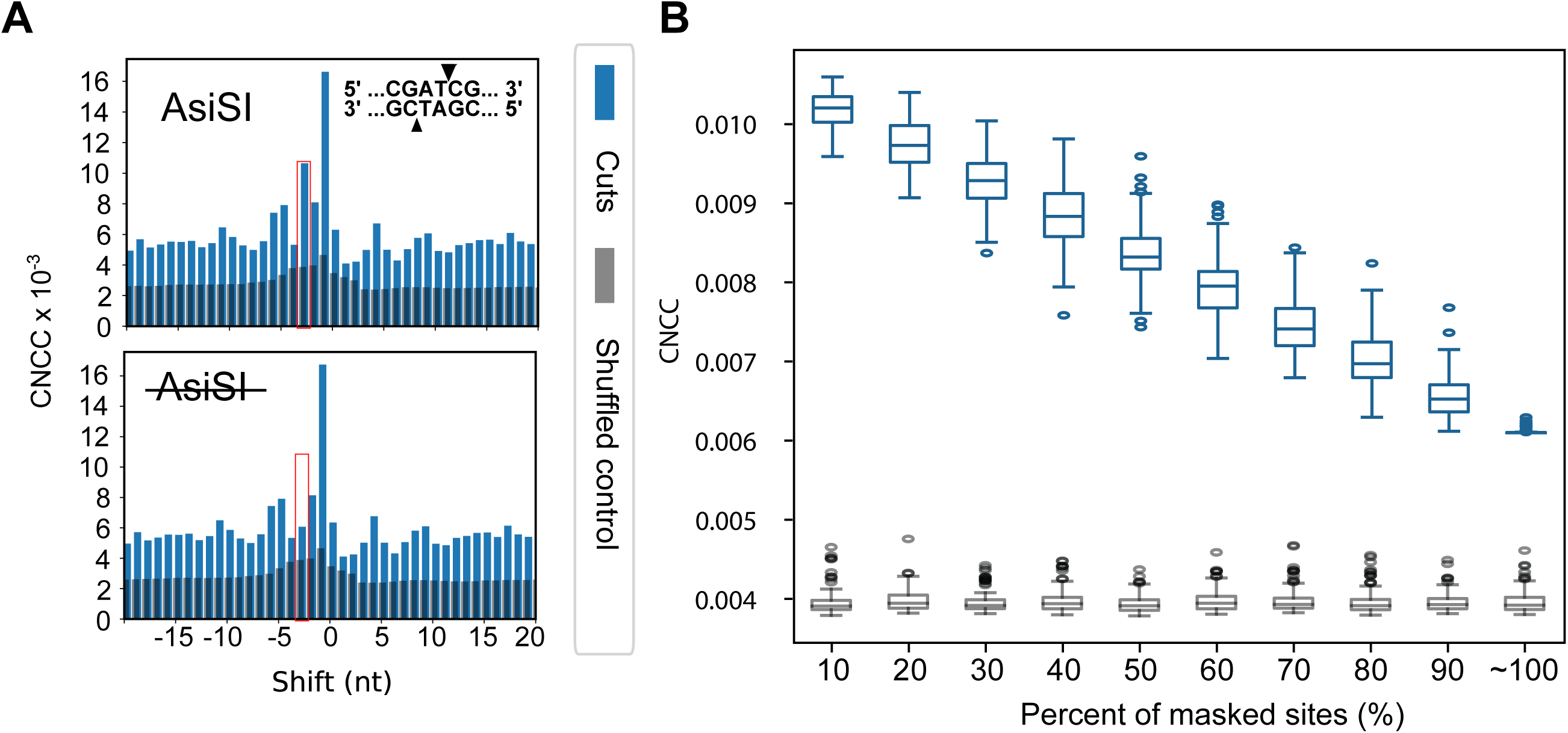
Analysis of CNCC end structure determination sensitivity. **(a)** CNCC analysis of DSBs generated by AsiSI limited digestion (top). When all AsiSI cut sites (0.03% of total reads) are masked, CNCC at −3 shift (marked by red rectangles) drops to the average level (bottom). (**b)** The CNCC signal at −3 shift (blue) with an increasing percentage of cut sites masked shows a concordant decrease in the CNCC signal at −3 shift, while the corresponding shuffled control remains constant (gray). Box represents 25% and 75%, and median is marked with the bar within the box. Whiskers depict 5% and 95%, and outliers are marked with ellipses.

### Investigating Endogenous and Etoposide-Induced Breaks

In the AsiSI analysis (**Fig. 2a**), along with the expected −3 spike, we observed a large break spike of CNCC at −1 shift, suggesting a high abundance of blunt end breaks. To determine if this spike was specific to U2OS cells or a generalizable endogenous species, we analyzed the CNCC break profile of untreated non-malignant lymphoblast cells, GM13069, and revealed a similar pattern (**Fig. S5a**). Further, endogenous breaks were enriched at promoters and transcription start sites (TSS), with 48% of the break density localized to these regions following read normalization for total region sizes (**Fig. S5b**).

This enrichment of reads at the TSS and promoter prompted us to examine the contribution of topoisomerase II (TOP II), which is known to act at promoters and TSSs [9-11]. To explore TOP II-mediated breaks, we treated GM13069 cells with three concentrations of etoposide, an inhibitor that prevents the ligation activity of TOP II, resulting in a covalently bound protein-DNA cleavage complex, which is known to be resolved by Mre11 [5, 12-14] or TDP2 [15-18]. Upon etoposide treatment, there was a dose-dependent increase in cell death examined by propidium iodide staining of the cells and flow cytometry analysis, to confirm the treatments (**Fig. S6**). The break mapping and sequencing was performed with two biological replicates of etoposide-treated samples. Pearson correlations between genome-wide read coverage of DSBs for each treatment showed a strong reproducibility between each biological replicate (r=0.86-0.98, **Fig. S7**). Further, there was a concordant and significant dose-dependent increase in break density at promoters and TSSs (p < 2.2×10^−16^, Kruskal-Wallis with Dunn test) (**Fig. 3a**), while all other regions lacked this increase, which is consistent with known locations of TOP II activity. This increase of breaks at TSSs and promoters following etoposide treatment has been previously suggested [19-21], and our genome-wide analysis directly confirms this change in break occurrence under etoposide treatment.

**Fig. 3.**
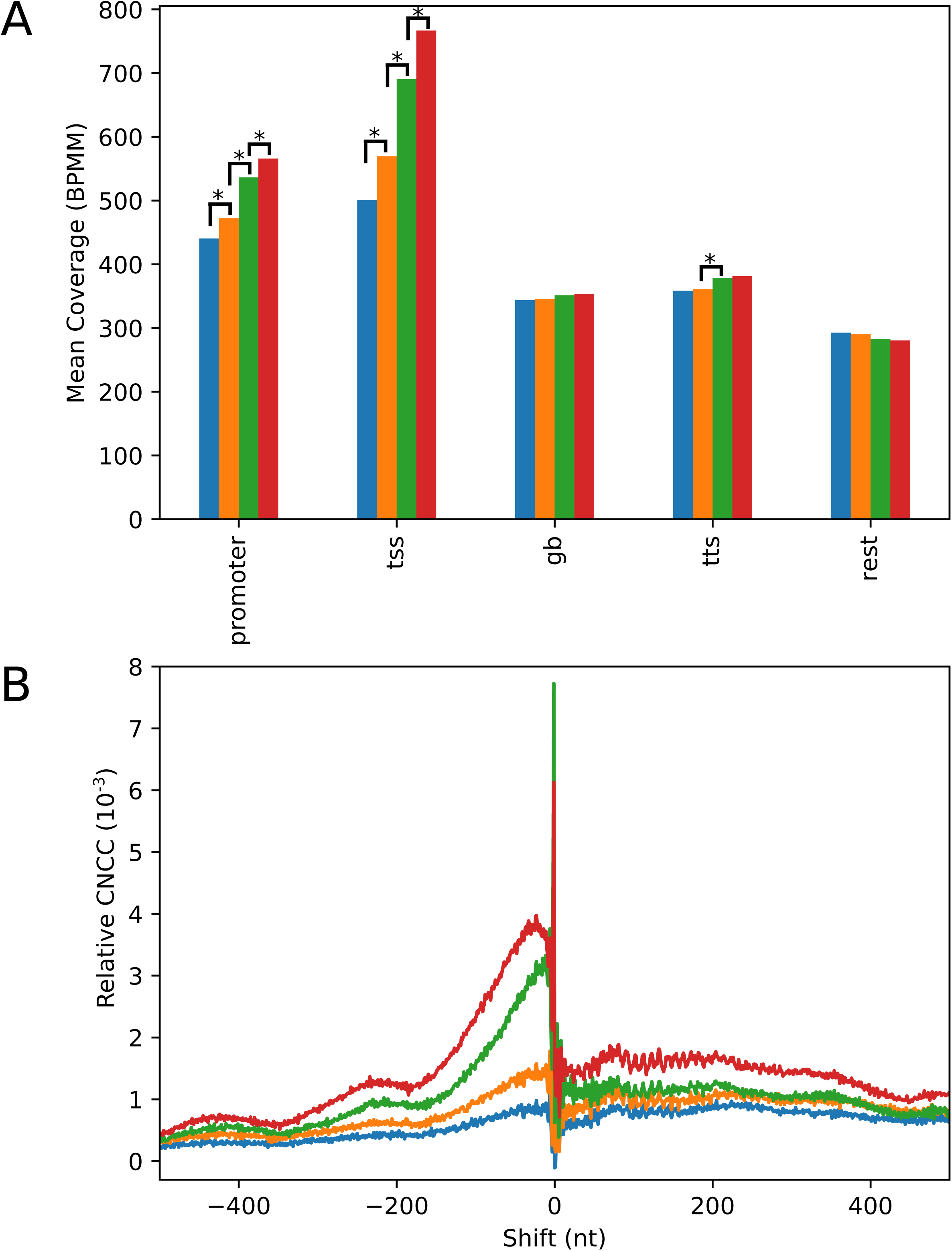
Inhibition of topoisomerase II with etoposide increases break densities at promoters and TSSs, reveals increased genome-wide 3’-overhang end structures, and displays the progression of 5’ to 3’ resection. **(a)** Total break density for two biological replicates in each annotated genomic region normalized to region size for all etoposide treatments (mean coverage as breaks per megabase per million (BPMM)). Increases in break density seen in the promoter and TSS are significant for each increasing etoposide treatment step (p < 2.2×10^−16^ is denoted as *). Significance was determined by Kruskal-Wallis test for each region, and followed up with Dunn tests using the Benjamini-Hochberg method of correction. **(b)** Treatment breaks of relative CNCC for etoposide-treated cells over a 1000-nt shift. Data represents the merge of two biological replicates for each treatment (**Table S2 and Fig. S9**).

Next, we applied the CNCC analysis to the two biological replicates of etoposide-treated samples separately (**Fig. S8**). Although there was slight variability between the replicates, the general shape and trend for each etoposide dose remained consistent, with the most variability being seen in the case of 15 μM which may be due to the high dosage. We also noticed that the CNCC analysis of the etoposide-treated samples displayed a much broader range of shifts. To establish an appropriate shuffled control for these samples, the 15 μM etoposide treated samples was used to test the same series of wiggle magnitudes as was done for HaeIII-digested sample (**Fig. S4b**). A wiggle magnitude of +2000 to −2000 (0, no shift, excluded) was identified to be meaningful and was implemented, because the maximum spike range to be controlled for is much larger (**Fig. S8**). We then merged the biological replicates, and performed CNCC on the merged data and generated respective shuffled controls for each treatment (**Fig. S9a**). To properly make comparisons between treatments, the median value of the shuffled control for each treatment was subtracted from the CNCC values of the corresponding treatment, to generate a “relative CNCC” (**Fig. 3b and Fig. S9b**). The relative CNCC signals of etoposide-treated samples showed that with increasing etoposide concentrations, there was an increase in the generation of CNCC spikes over a broad range of negative shifts (**Fig. 3b**). This broad range of spikes which have shift values less than −1 indicates the generation of 3’ overhang ends, and the shape of the CNCC spikes suggests a resection gradient, with a maximum length of approximately 165 nts. This 5’ to 3’ resection is likely the result of TOP II cleavage complexes being removed and processed by Mre11 or TDP2. TOP II does not have precise recognition sequences [9, 22, 23], therefore, the display of the 5’ to 3’ end resection globally by the CNCC analysis without knowing specific targeted sequences demonstrates the valuable utility of CNCC. Moreover, the 1.5 μM treatment revealed a peak of CNCC resection signature between 4 – 28 nts (determined by 90% of maximum CNCC value in the resection range), representing the most common resected 3’ overhang end structure following this dose of etoposide treatment. The peak of the CNCC resection signature observed for 15 μM etoposide treatment was 5 – 45 nts, demonstrating that the length of resection increased with higher etoposide treatment and that our analysis is capable of distinguishing even these fine differences in the end structure distributions between treatments.

## DISCUSSION

While cross correlation analyses have been previously applied to ChIP-seq data, this is the first time this analysis approach has been applied to genome-wide break mapping/sequencing data. We took advantage of the ability of cross correlation analysis to identify patterns in a noisy background, and combined this with the single nucleotide resolution of the break mapping/sequencing data to analyze structures following DNA damage. We demonstrated the ability for our method of coverage-normalized cross correlation analysis to determine the genome-wide end-structure distribution of DNA double-stranded breaks at single-nucleotide resolution. Our analysis tool has proven to work for both induced break systems (sequence-specific breaks by restriction enzymes and etoposide-induced breaks) to capture the resultant break end structure landscape of the cell. For the first time on a genome-wide scale, our method revealed the increase in the 5’ to 3’ end resection following etoposide treatment, and more importantly, the global progression of the resection due to the removal of DNA topoisomerase II cleavage complexes. The change in extent of resection could indicate a change in which pathways are being used to repair the DSBs. The difference in DNA end structure at the site of break and the extent of resection in part dictates repair pathway choice between NHEJ, HR, and other pathways such as microhomology-mediated end joining (MMEJ) and single-strand annealing (SSA) [2, 3, 24-27]. While little to no resection and end-processing is supportive of NHEJ, short resection facilitates a shift towards SSA and MMEJ, but a long resection drives completely to HR. Further investigation into different repair pathways, or specific proteins in these pathways, can be benefited from including mapping genome-wide breaks and coupling with our CNCC analysis to determine the impacts on the repair of various endogenous and induced breaks.

In addition to the ability to determine end structures at DSBs, the CNCC analysis of broken ends allows for identification of consensus sequences (if they exist) located at the breaks. Using the break data from the BanII digestion, we identified pairs of DSB coverage containing at least two reads, which displayed the spike on the negative strand located 4 nts upstream from the spike on the positive strand (n = 718,163) (example genomic sites, **Fig. S10a**). By performing a motif analysis of these read pair sites, we can recapitulate the BanII consensus sequence (**Fig. S10b**). This analysis demonstrates that the knowledge of end structure from CNCC analysis can be further used to understand potential sequence motifs associated with identified end structures.

The ability of CNCC to determine DNA break end structures without *a priori* knowledge of the break sequences or genomic locations can lend itself to multiple analyses, provided the end-structure or break species under study is appropriately sampled in the sequencing data. Our method can be applied to genome-wide DSB sequence mapping datasets over a broad range of treatment conditions and across cell types to better understand the impact and specificity of treatments on generating breaks, and to investigate the fate of broken ends and proteins that repair them.

## MATERIALS AND METHODS

### Cell culture and treatments

GM13069 cells, a human lymphoid cell line derived from a normal individual (Coriell Institute), were grown in RPMI 1640 medium (Gibco 11875) supplemented with 10% fetal bovine serum and plated at 2 × 10^6^ cells per 100mm cell culture dish. Cells were treated 18 hours later with etoposide (Sigma E1383) at 0.15 μM, 1.5 μM, or 15 μM for 24 hours. Cells and cell culture medium were then collected by centrifugation at 4 °C and washed twice with cold PBS containing the treatment dose of etoposide. After washes, cells were divided into two equal aliquots. For cell viability assay, one aliquot of cells was resuspended in 2 ng/mL propidium iodide for flow cytometry analysis using a FACSCalibur flow cytometer (BD Biosciences). For breakpoint detection, genomic DNA was purified from the other aliquot of cells by lysing cells in 50mM Tris.Cl (pH8.0); 100mM EDTA; 100mM NaCl; 1% SDS; 1mg/mL Proteinase K for 3 hours at 55 °C followed by organic extraction purification and ethanol precipitation. Precaution such as gentle pipetting with wide-opening pipette tips to avoid shredding DNA was taken to avoid introduction of DNA breaks during purification.

### Genome-wide break mapping and sequencing

Detection of DSBs was performed as described [8] with modifications. Briefly, genomic DNA from etoposide treated cells or restriction enzyme-digested DNA were subjected to end-blunting reactions with T4 DNA polymerase, Klenow fragment of DNA Polymerase I, and T4 Polynucleotide kinase. During the reaction (Fig. 1), two blunted ends of each break will stay as blunted; two break ends with 3’ overhangs are trimmed by the 3’ to 5’ exonuclease activity of the polymerases to generate two blunted ends with a separation based on the reference sequence; two break ends with 5’ overhangs are filled-in by the 5’ to 3’ polymerase activity of the polymerases to generate two blunted ends with a overlap based on the reference sequence. The CNCC analysis utilizes the separation or overlap of the two ends to distinguish these three end structures. The end-blunting reactions are followed by A-tailing reactions and Illumina adaptor P5 ligation to broken DNA ends. Excess adaptor was removed and then DNA was fragmented by sonication, and subsequently ligated to Illumina adaptor P7. The libraries were amplified by PCR for 15 cycles. Prepared libraries were then subjected to whole-genome 75 bp paired-end sequencing by the Illumina NextSeq 500 platform.

### Sequencing data analysis

Sequencing reads were aligned to the human genome (GRCh38/hg38) with bowtie2 (v.2.3.0) aligner running in high sensitivity mode (*--very-sensitive*, critical program options are given in parentheses). Restriction on the fragment length from 100 nt to 2000 nt (*-X 2000 -I 100* options) was imposed. Unmapped, non-primary, supplementary and low-quality reads were filtered out with samtools view (v. 1.7) (*-F 2820*). Further, to ensure reads from independent events, PCR duplicates were marked with picard-tools (v. 1.95) MarkDuplicates and finally, read 1s (*-f 67*) from non-duplicated reads (*-F 1024*) were filtered with samtools view for continued analysis. Further downstream analysis was performed with BEDtools (v. 2.26.0). Data were visualized in python (v. 3.5.2) with libraries: pandas (v. 0.22.0), numpy (v. 1.13.3), and matplotlib.pyplot (v. 2.0.2).

### Coverage-Normalized Cross Correlation

First, genome-wide coverage was calculated with bedtools genomecov for each strand separately (*-strand +/-*) using only the 5’ end (*-5* option) of read 1 of non-duplicated reads (see aligning procedure described above). Output files were saved in dz format (*-dz* option). These coverage output files were then read into Jupyter notebooks with python (v. 3.5.2) and used pandas (v. 0.22.0) data frames to implement coverage-normalized cross correlation calculations (with numpy.dot function), at which time coverage over the centromeres was masked (discarded) prior to final calculation.

Coverage-normalized cross correlation (CNCC) is a cross correlation that is further normalized based on overall coverage as to allow for meaningful comparison between different biological samples.

CNCC as such is defined as:

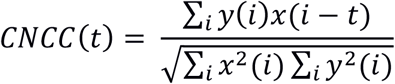

where: *x(i)* is a DNA break coverage on the positive strand at position *i, y(i)* is a DNA break coverage on the negative strand at position *i*, and *t* is the shift distance of interest. (Here we are using only the most 5’ nt of read 1 as it exactly maps DNA break position).

Modified, two-step shuffled control was calculated by using the coverage output files from above then implementing both a wiggle of position using random wiggle assignment (with numpy.random.randint) and a shuffling of the coverage values between positions (with numpy.random.permutation) for each sample. Then, CNCC, as defined above, was calculated for the minimally perturbed data to generate the stringent shuffled control. For the restriction enzyme digested samples, we applied a wiggle value ranging from +2 to −2 (0 excluded). For the etoposide-treated samples, we applied a wiggle value ranging from +2000 to −2000 (0 excluded) (see main text for the discussion on these shuffled controls).

### Sensitivity of CNCC analysis

Published break mapping data from a limited AsiSI digestion in U2OS cells [8] was aligned and processed by the CNCC analysis. The AsiSI enzyme was fused to an estrogen receptor ligand binding domain, and when cells were treated with 4-hydroxytamoxifen, the enzyme was transported to the nucleus where it then generated DSBs at loci containing its consensus sequence. This resulted in a limited digestion where reads from AsiSI cleavage (mapped to the consensus sequence) were only 0.03% of total reads. CNCC analysis was first implemented on the complete data set, and then on the data set that filtered out all reads mapped to AsiSI locations, to assess the ability of CNCC to detect change. To further test sensitivity, AsiSI cut sites were masked in increments of 10%, CNCC was performed as above, and output for only the −3 shift position (the AsiSI-induced spike) was evaluated. For each 10% increment, there were 100 iterations of site masking, and only sites with more than 5 reads at the site were included in the analysis (n = 303). In each iteration for a 10% increment, the number of randomly picked AsiSI cut sites remained constant, while the number of reads that were masked varied as a result.

### BanII consensus sequence analysis

Pairs of DSB spikes separated by 4 nt (negative strand spike upstream from spike on positive strand) were found. Of those pairs, 99.5% (n = 714,922) were within canonical cut sites of BanII in the reference sequence, and the remaining 0.5% of pairs (n = 3241) contain sequences that differ by only 1 nt from the canonical enzyme preference sequence. Sequences at all of these loci were extracted with BEDtools getfasta and the motif was found by DREAM from MEME Suit (v. 4,12.0).

### Statistical Analysis

For the comparison of genomic coverage between technical duplicates of BbvI digestion and between each biological duplicate for etoposide treatment, Pearson’s correlation was calculated for the genome-wide coverage in 100-nt and 1000-nt non-overlapping windows, respectively. For the dose-dependent change in break density following etoposide treatment, Kruskal-Wallis test was performed in R (v. 3.4.3) on the normalized coverage data in each genomic region. Significant results from Kruskal-Wallis were then followed up with a Dunn test, using the Benjamini-Hochberg method, to then determine the significance of the changes between each treatment level for the specific region.

**Abbreviations used**
DSB: DNA double-stranded break
HR: homologous recombination
NHEJ: non-homologous end joining
CNCC: coverage-normalized cross correlation
nt: nucleotide
TSS: transcription start sites
TOP II: topoisomerase II
MMEJ: microhomology-mediated end joining
SSA: single-strand annealing

## Data Availability

Data generated and reported in Fig. 1b, Fig. 3, Tables S1-2, and Figs. S2, S4-9 (BbvI, BanII and HeaIII digestion, and TOPII inhibition) has been deposited in Sequence Read Archive under the accession number: PRJNA497476 (https://www.ncbi.nlm.nih.gov/sra/PRJNA497476).

Publicly available data used in this study can be accessed in GEO under accession number: GSE78172 (EcoRV and AsiSI digestion experiments).

## Acknowledgements

The authors would like to thank Kristyna Kupkova for her initial suggestion of using cross correlation analysis and Sandeep Singh for critical reading of the manuscript. H.M.R. was supported by the Cellular and Molecular Biology Predoctoral Training Grant from the NIH (T32GM008136). This work was supported by grants from NIGMS (RO1GM101192) and NCI (RO1CA113863) to Y.H.W.

## Supplementary data

Supplementary Data to this article can be found online.

