## Supplemental Data for "CNCC: An analysis tool to determine genome-wide DNA break end structure at single-nucleotide resolution"

**Supplementary Data**

Table S1. Reads and cut sites statistics of restriction enzyme-digested samples

Table S2. Sequencing and alignment statistics

Figure S1. Outline of genome-wide break mapping protocol.

Figure S2. Reproducibility of genome-wide break mapping in BbvI digestion experiment.

Figure S3. Coverage-normalized cross correlation for EcoRV-digested DNA from HeLa cells.

Figure S4. Determination of appropriate control for coverage-normalized cross correlation.

Figure S5. Coverage-normalized cross correlation calculation for endogenous breaks and annotation of locations for all breaks.

Figure S6. Evaluation of etoposide treatment on cell survival.

Figure S7. Reproducibility of genome-wide break mapping for different etoposide treatments in GM13069.

Figure S8. Coverage-normalized cross correlation of breaks upon treatment maintains similar trend between biological replicates for GM13069.

Figure S9. Coverage-normalized cross correlation calculated for breaks upon treatment for combined biological replicates showing shuffled control and relative CNCC signal.

Figure S10. Determination of consensus sequences at enzyme cut sites leveraging coverage-normalized cross correlation.

**Table S1.** Reads and cut sites statistics of restriction enzyme-digested samples

| Enzyme | # of RE* cut sites with reads | Median read count per RE cut site | Total mapped end reads | Reads at RE cut sites (% of total reads) |
|---|---|---|---|---|
| Ban II | 2,064,247 | 6 | 15,046,243 | 13,546,297 (90%) |
| Bbv I | 2,331,244 | 4 | 31,754,252 | 19,374,772 (61%) |
| Hae III | 3,926,902 | 2 | 16,614,091 | 16,074,221 (97%) |

*RE: restriction enzyme

**Table S2.** Sequencing and alignment statistics

| Cell type | Treatment | Replicate | | Sequenced read pairs | Alignment rate | Duplication rate | Mapped DNA ends |
|---|---|---|---|---|---|---|---|
| | | Biological | Technical | | | | |
| GM13069 | BbvI | 1 | 1 | 16,442,364 | 94.81% | 6.09% | 11,426,508 |
| | | | 2 | 30,598,266 | 94.14% | 8.42% | 20,327,744 |
| | BanII | 1 | 1 | 21,575,010 | 95.09% | 6.32% | 15,046,243 |
| | HaeIII | 1 | 1 | 25,733,886 | 93.32% | 5.31% | 16,614,091 |
| | UT | 1 | 1 | 11,226,462 | 82,18% | 16.64% | 6,069,143 |
| | | | 2 | 20,763,138 | 68.90% | 22.38% | 8,729,170 |
| | | 2 | 1 | 30,265,469 | 90.09% | 31.49% | 15,634,772 |
| | Etoposide 0.15 µM | 1 | 1 | 19,176,118 | 70.18% | 26.98% | 7,871,510 |
| | | | 2 | 10,502,446 | 96.12% | 11.80% | 7, 371,345 |
| | | 2 | 1 | 26,165,279 | 93.56% | 25.91% | 14,601,975 |
| | Etoposide 1.5 µM | 1 | 1 | 12,894,905 | 92.45% | 13.09% | 8,552,854 |
| | | | 2 | 22,516,349 | 88.45% | 18.62% | 13,355,567 |
| | | 2 | 1 | 26,509,305 | 94.64% | 11.01% | 18,191,095 |
| | Etoposide 15 µM | 1 | 1 | 14,522,657 | 94.38% | 10.86% | 9,803,647 |
| | | | 2 | 19,049,613 | 92.12% | 12.83% | 12,295,170 |
| | | 2 | 1 | 32,301,001 | 92.11% | 12.86% | 21,699,825 |

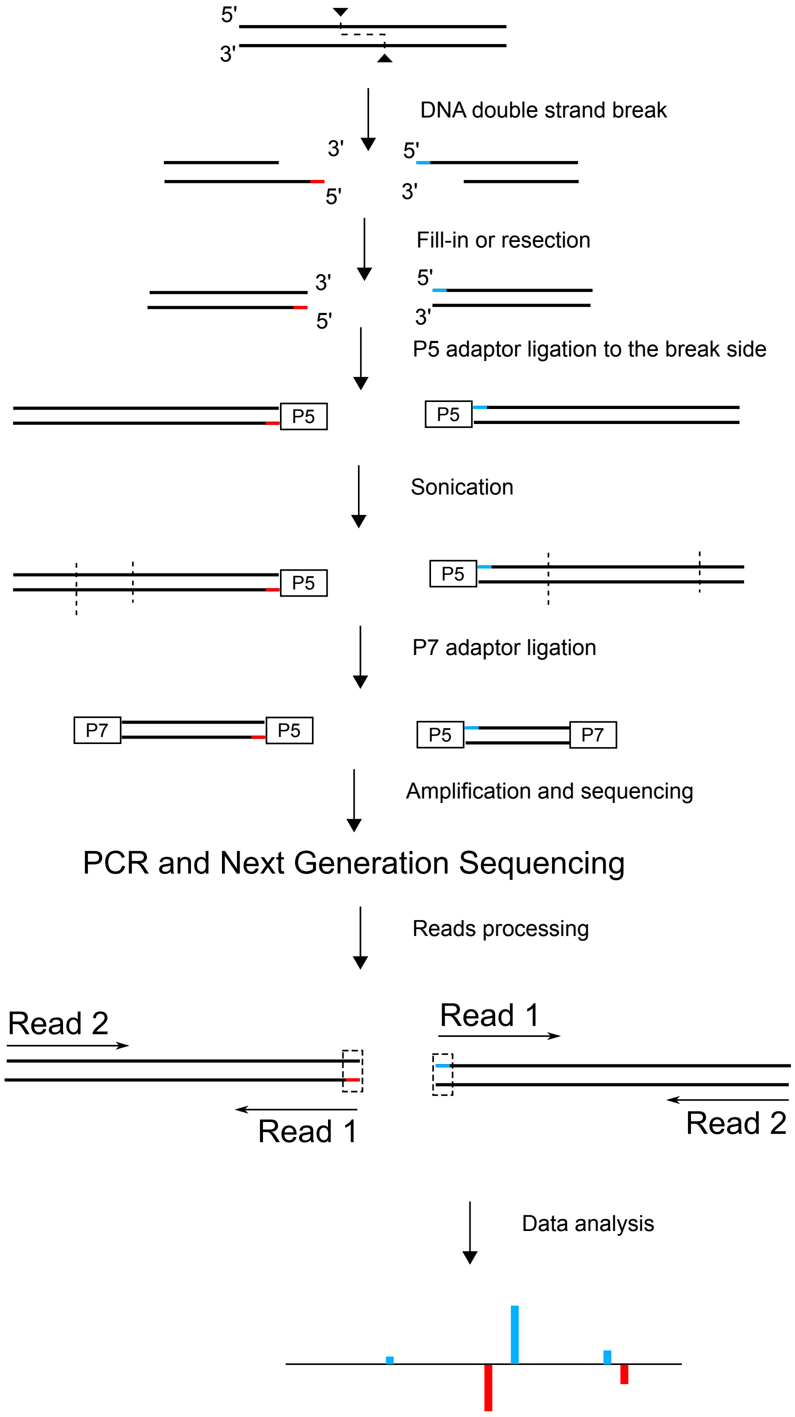


**Figure S1. Outline of genome-wide break mapping protocol.** Every double-stranded DNA break generates two DNA ends (indicated in blue and red). These ends are then either filled-in (5' overhang) or resected (3' overhang) to produce blunt ends. The critical step is to ligate the P5 adaptor, effectively capturing the 5’ end of the break for each of the two ends and then removing excess P5 adaptor before sonication. Next, the P7 adaptor is ligated to the artificially generated breaks from sonication. As a result of the precisely controlled adaptor ligation, only fragments carrying information about double-strand breaks are amplified and sequenced because only such fragments have both a P5 and P7 adaptor. The use of paired-end sequencing then allows for meaningful removal of PCR duplicates from the data. For further analysis we extract only the first nucleotide of the read 1s as they precisely map the DNA break locations as outlined in **Fig. 1**.

**(a)**

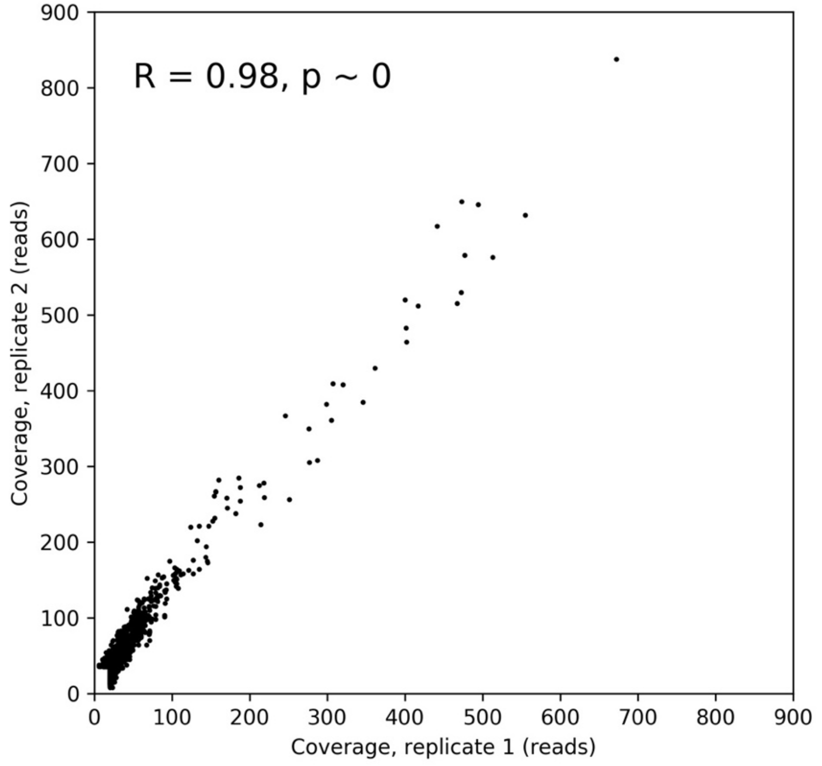


**(b)**

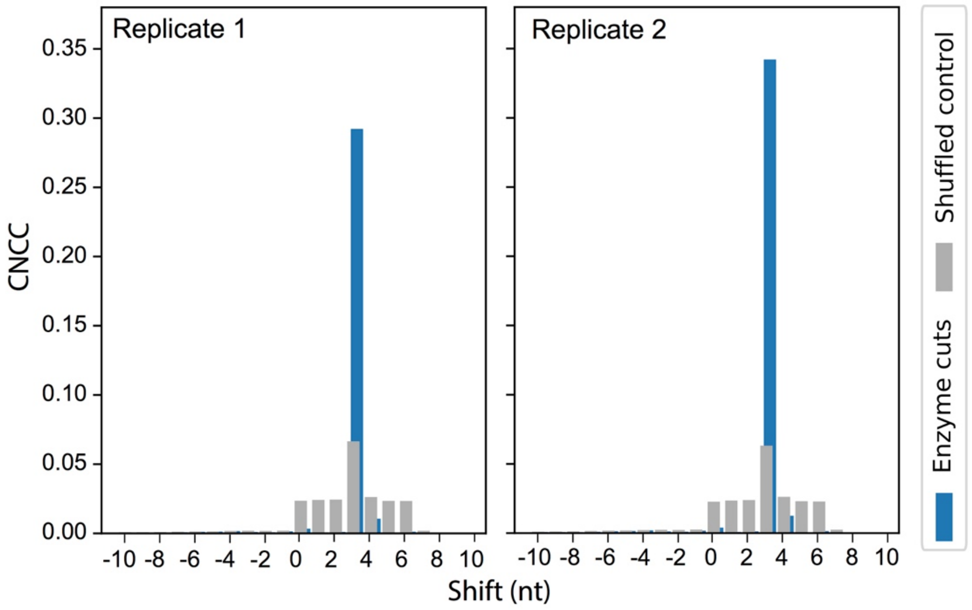


**Figure S2. Reproducibility of genome-wide break mapping in BbvI digestion experiment. (a)** Scatter plot presenting genome-wide coverage of the genome-wide break mapping reads calculated for 100-nt non-overlapping windows. Windows with reads less than 10 in both replicates are excluded from this analysis. Pearson correlation coefficient and p-value are shown. **(b)** The same, strong CNCC spikes at the shift of +3 (blue bar) were observed for both replicates calculated separately. To create a stringent shuffled control (gray), we introduced small perturbations to the signal to conserve the overall composition of the break intensities and clustering (±2-nt wiggle, see "Materials and Methods" for details).

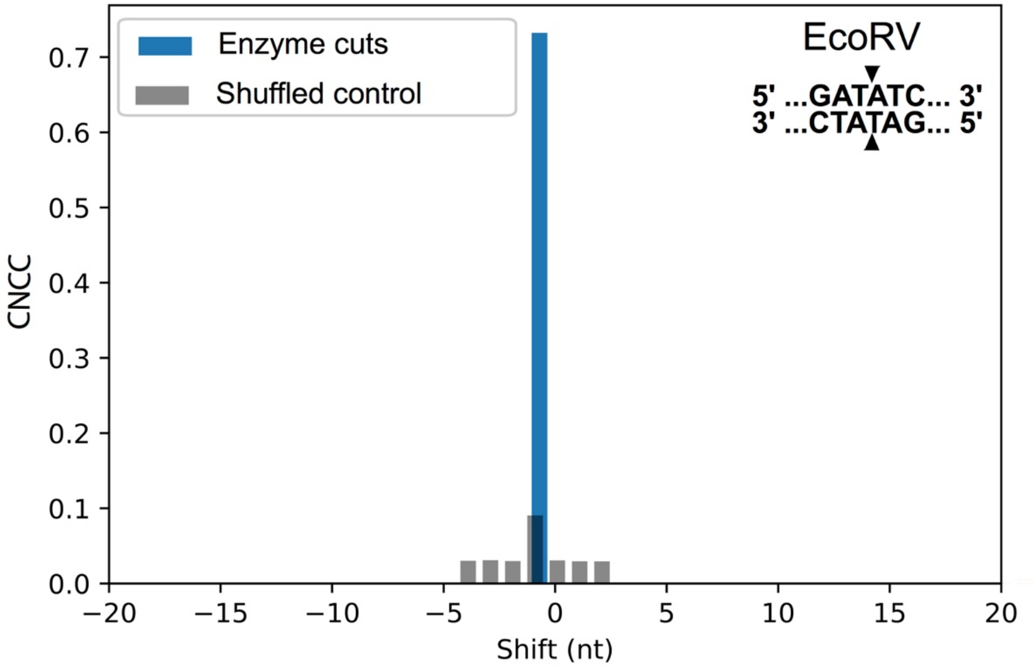


**Figure S3. Coverage-normalized cross correlation for EcoRV-digested DNA from HeLa cells.** The restriction enzyme EcoRV cuts DNA producing blunt ended breaks, as indicated in the upper right. As shown in **Fig. 1a,** break results in a strong CNCC spike at -1 (blue bar).

(a)

(b)

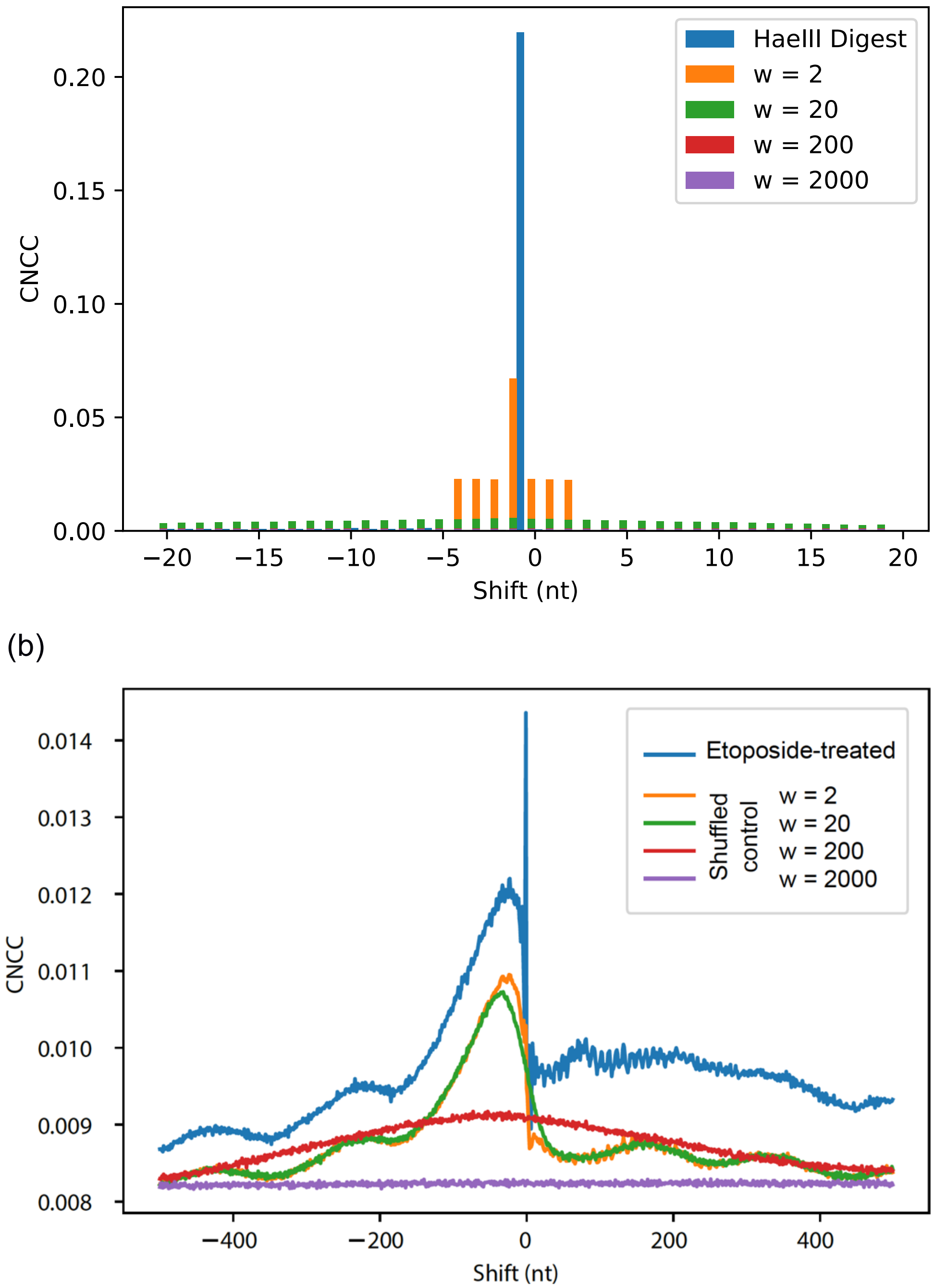


**Figure S4. Determination of appropriate control for coverage-normalized cross correlation.** CNCC values are shown for breaks generated from Hae III-digested DNA **(a)** or DNA from 15 µM etoposide-treated GM13069 cells (combined biological replicates 1 and 2) **(b)**. Different perturbations of the shuffled control are shown with different ranges of wiggled positions (w). The magnitude of the wiggle directly corresponds to the maximum spike range the control is needed for. Ultimately, for the restriction enzyme digested samples, we applied a wiggle ranging from +2 to -2 (0 excluded). For the etoposide-treated samples, we applied a wiggle ranging from +2000 to -2000 (0 excluded).

(a)

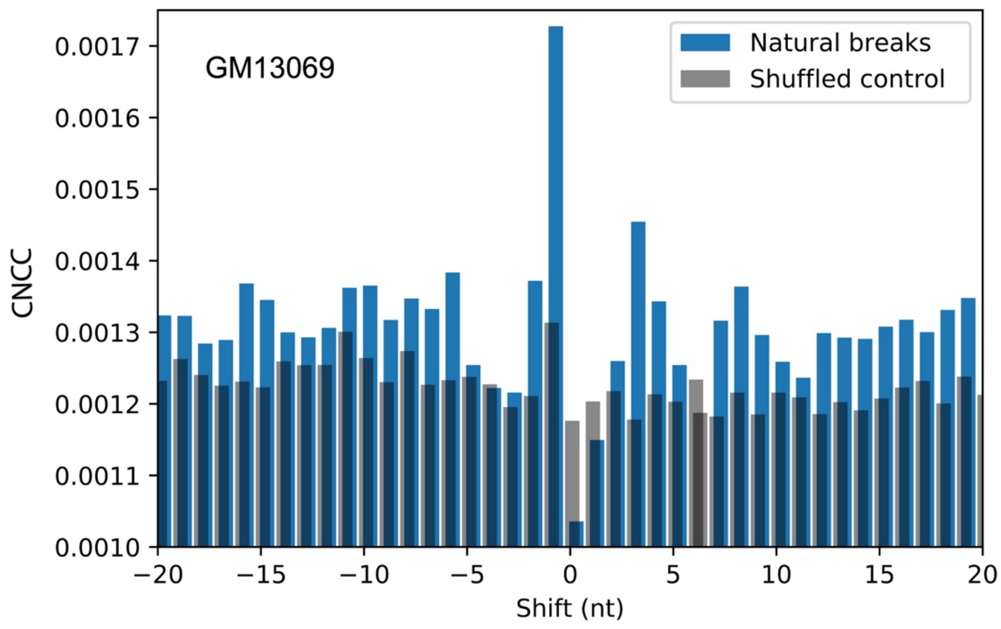


(b)

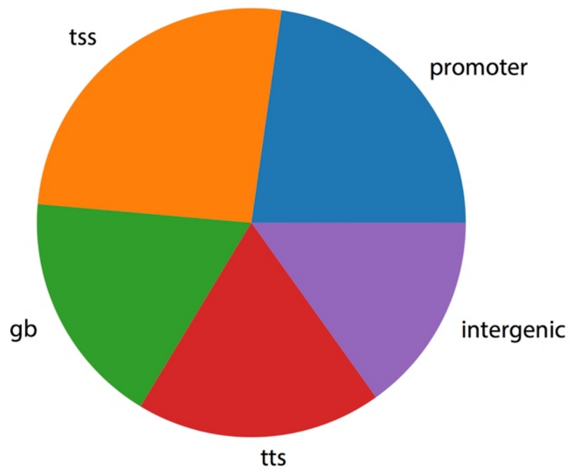


**Figure S5. Coverage-normalized cross correlation calculation for endogenous breaks and annotation of locations for all breaks. (a)** CNCC calculated for untreated GM13069 lymphoblast cells indicates various shift positions, with natural breaks correlation above that of shuffled control. **(b)** Annotation of all read sites indicates 48% of all break density (number of reads adjusted to the total region size) is in transcription start sites (tss), and promoters, while the rest distributes among gene bodies (gb), transcription termination sites (tts), and intergenic regions.

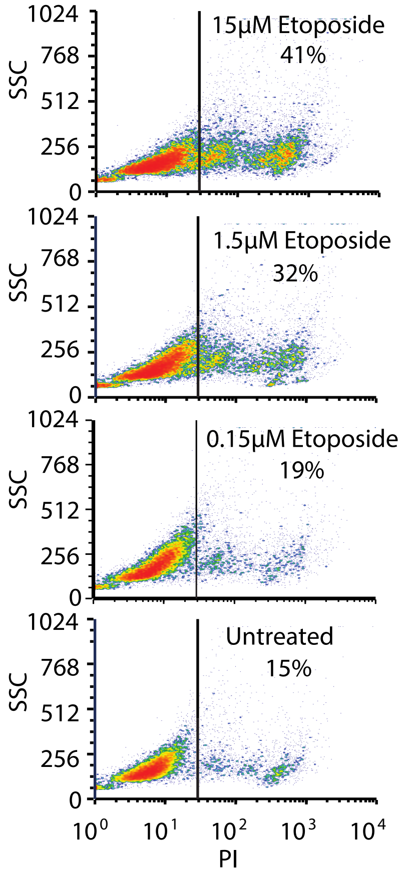


**Figure 6. Evaluation of etoposide treatment on cell survival.** Flow cytometry analysis of propidium iodide (PI)-stained GM13069 cells for each treatment shows the expected increased cell death following increased etoposide concentration (data represents biological replicate 1).

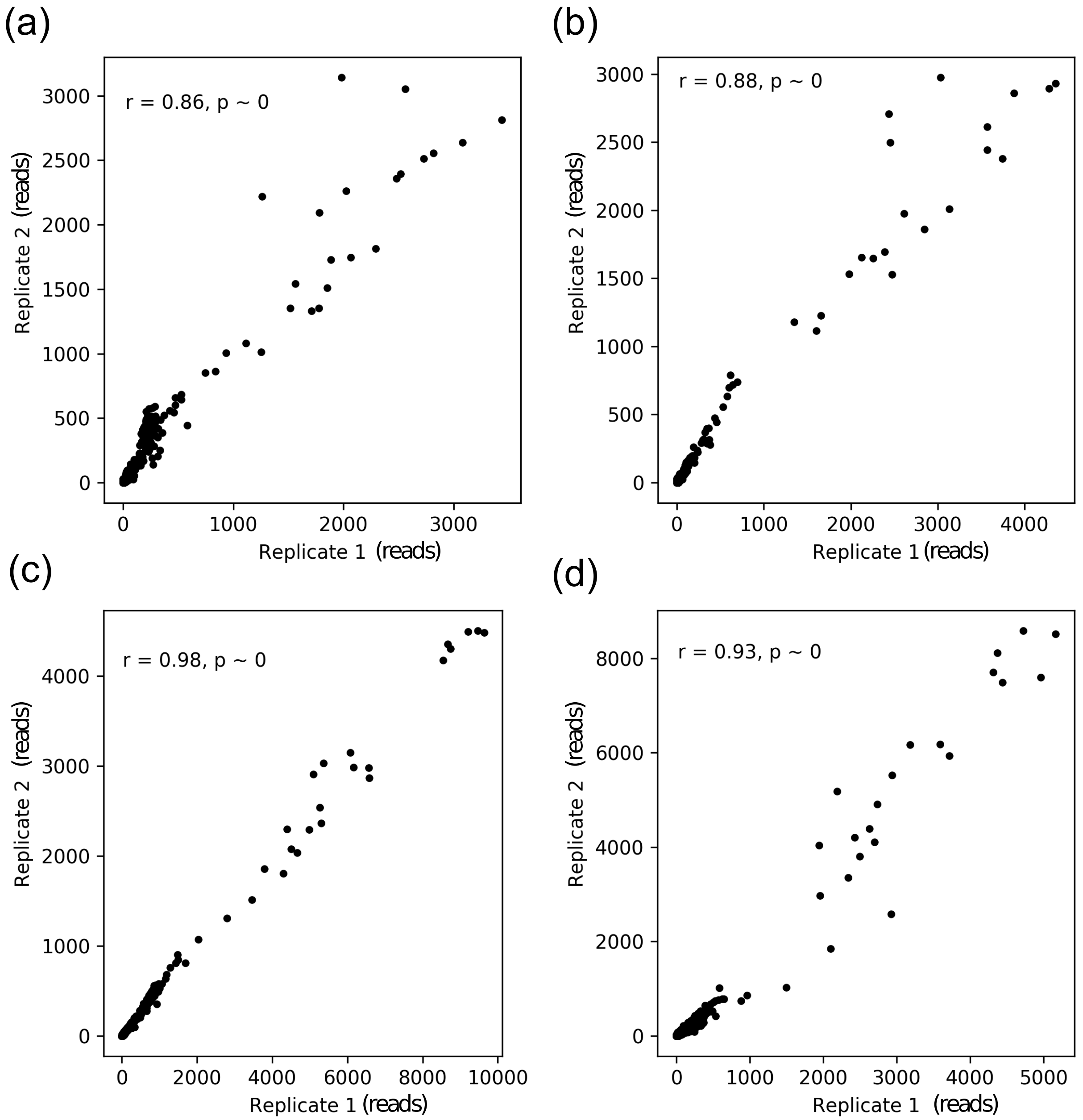


**Figure S7. Reproducibility of genome-wide break mapping for different etoposide treatments in GM13069**. Scatter plots presenting genome-wide coverage of the genome-wide break mapping reads calculated for 1000-nt non-overlapping windows for two biological replicates. Windows with no reads in both replicates are excluded from analysis. Pearson correlation coefficient and p values are shown for untreated **(a)**, 0.15 μM **(b)**, 1.5 μM **(c)**, and 15 μM **(d)** etoposide-treated samples. Sequencing and read information can be found in **Table S2**.

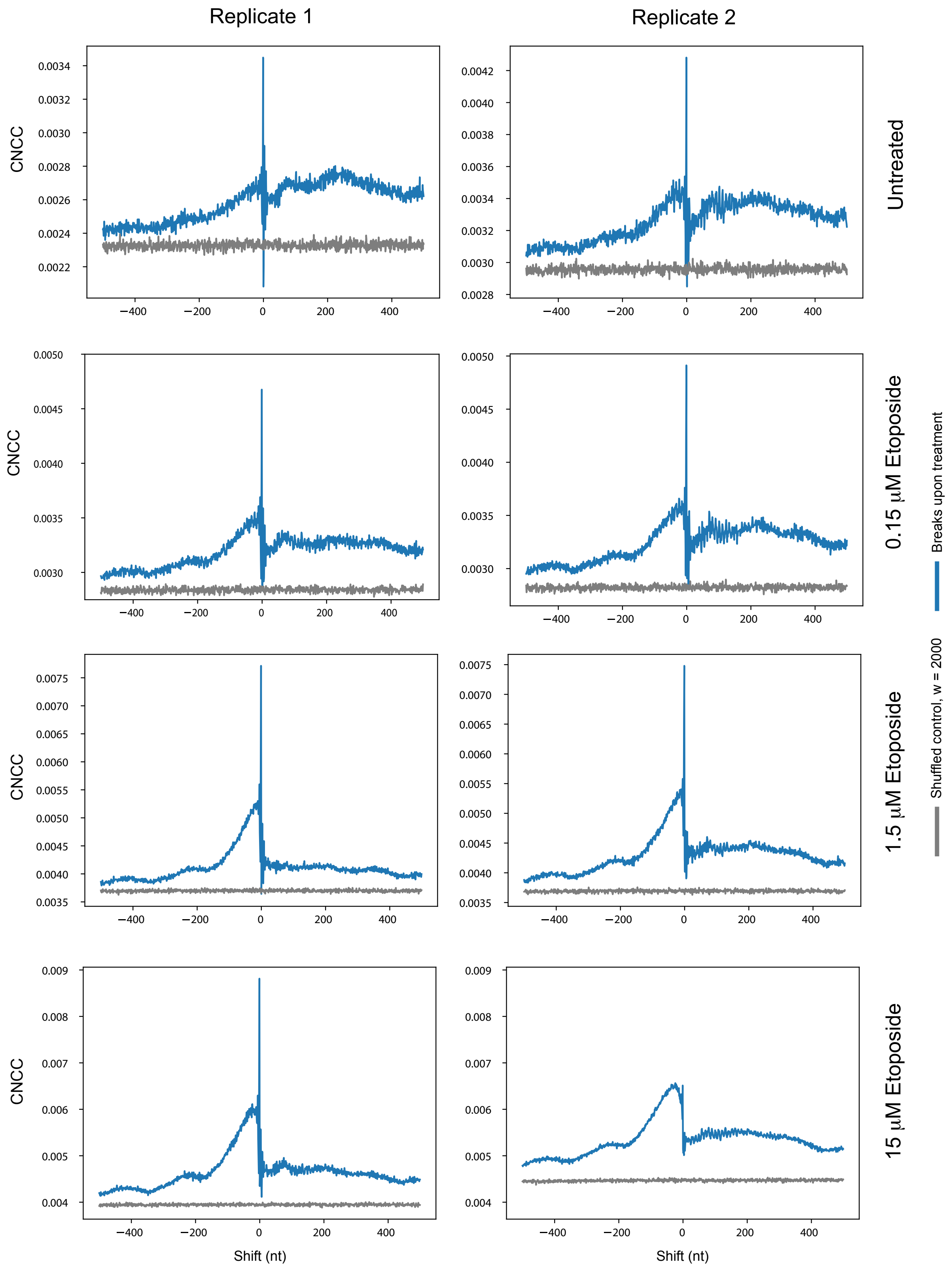


**Figure S8. Coverage-normalized cross correlation of breaks upon treatment maintains similar trend between biological replicates for GM13069.** Side-by-side comparison of the calculated CNCC for the breaks captured upon treatment of the two biological replicates for each etoposide treatment. Overwhelmingly, the signal compared between replicates is consistent, with the only slight divergence being found in the 15 μM etoposide treatment.

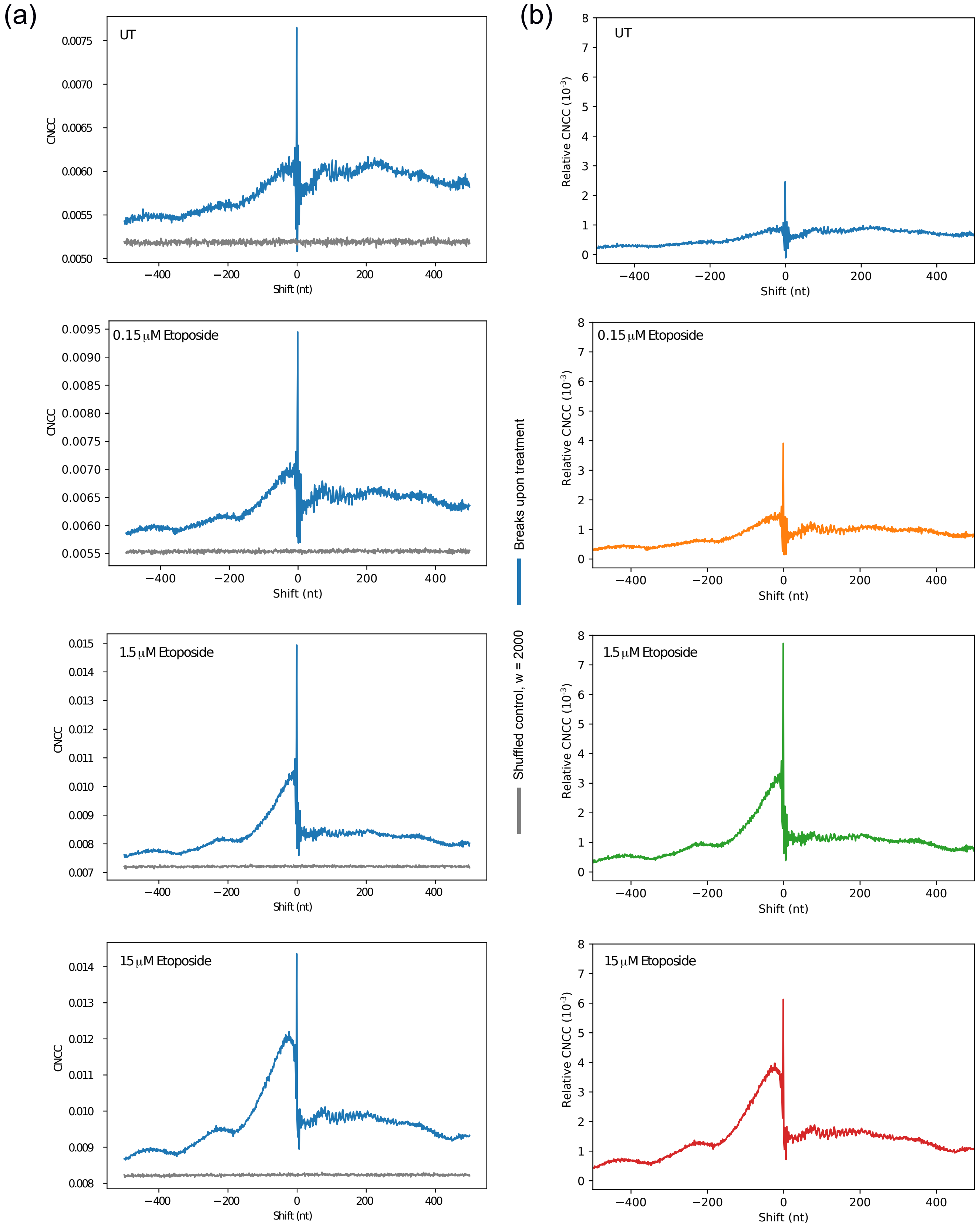


**Figure S9. Coverage-normalized cross correlation calculated for breaks upon treatment for combined biological replicates showing shuffled control and relative CNCC signal. (a)** CNCC signals (blue) shown for breaks upon etoposide treatments of GM13069 combining two biological replicates. The median value for the shuffled control (gray) was subtracted from the treatment breaks signal to make each respective relative CNCC signal **(b)** also shown in **Fig. 3b**.

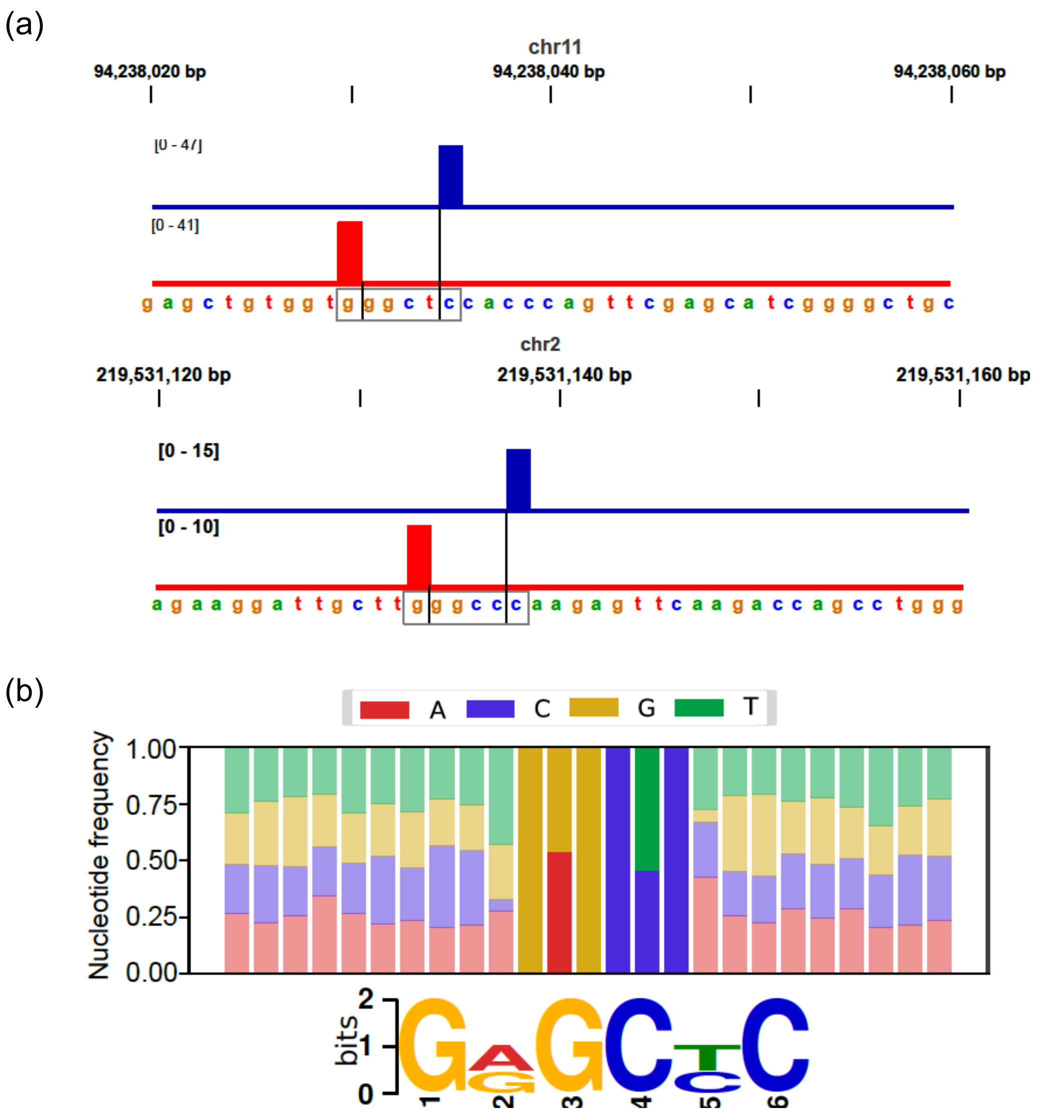


**Figure S10. Determination of consensus sequences at enzyme cut sites leveraging coverage-normalized cross correlation.** As shown on **Fig. 1b** cross correlation suggests that the majority of cuts from BanII digestion produce reads that are separated by 5 nt (representing a 4-nt 3' overhang). **(a)** Survey of such read pairs (two examples are shown) revealed 721,403 potential cut sites. Coverage of the 5' ends of genome-wide break mapping reads on the positive (blue) and negative (red) strands demonstrates that these reads correspond to enzyme cut sites as indicated by the nucleotide sequences shown below each coverage profile matching the BanII recognition site sequence (GRGCYC). **(b)** Analysis of all the potential site sequences perfectly recapitulates the enzyme consensus sequence in both: nucleotide frequency (top) and motif search analysis (bottom). Motif was found by DREAM from MEME Suit (v. 4,12.0) with e-value $1.3\text{x}10^{-30873}$. For comparison, the next found motif's e-value was $1.7\text{x}10^{-1275}$.
